# Consistent hierarchies of single-neuron timescales in mice, macaques and humans

**DOI:** 10.1101/2024.10.30.621133

**Authors:** Zachary R. Zeisler, Marques Love, Ueli Rutishauser, Frederic M. Stoll, Peter H. Rudebeck

## Abstract

The intrinsic timescales of single neurons are thought to be hierarchically organized across the cortex. This conclusion, however, is primarily based on analyses of neural responses from macaques. Whether hierarchical variation in timescales is a general brain organizing principle across mammals remains unclear. Here we took a cross-species approach and estimated neuronal timescales of thousands of single neurons recorded across multiple areas in mice, monkeys, and humans using a task-agnostic method. We identify largely consistent hierarchies of timescales in frontal and limbic regions across species: hippocampus had the shortest timescale whereas anterior cingulate cortex had the longest. Within this scheme, variability across species was found, most notably in amygdala and orbitofrontal cortex. We show that variation in timescales is not simply related to differences in spiking statistics nor the result of cytoarchitectonic features such as cortical granularity. Thus, hierarchically organized timescales are a consistent organizing principle across species and appear to be related to a combination of intrinsic and extrinsic factors.

**Highlights:**

- Similar hierarchies of single neuron timescales exist across humans, macaques, and mice
- Humans diverge from an otherwise conserved hierarchy in amygdala and OFC
- Timescale variability in ventral frontal cortex is not entirely related to cytoarchitecture

## Introduction

Receptive fields are a foundational concept in psychophysics and neuroscience, a classic example being the spatial receptive fields of single neurons in primary visual cortex (Hubel and Wiesel, 1962; Spillmann, 2014) and higher-order parietal and frontal areas (Viswanathan and Nieder, 2017). An influential line of work has extended this framework to the temporal domain (Murray et al., 2014). Using the rate of exponential decay of the spike-count autocorrelation, Murray and colleagues extracted the intrinsic temporal receptive field, or timescale, of a single neuron or population of neurons. These neural timescales, which are hypothesized to reflect the amount of time over which a neuron integrates information, appear to be hierarchically organized across the brain (Murray et al., 2014; Nougaret et al., 2021). Neurons within sensory cortex have shorter timescales, while neurons within higher-order associative areas, particularly in frontal cortex, have longer timescales. Convergent evidence from fMRI in humans (Hasson et al., 2008; Honey et al., 2012; Raut et al., 2020), fMRI and neurophysiology recordings in non-human primates (Murray et al., 2014; Manea et al., 2022, 2024), and modelling work (Gao et al., 2020; Perez-Nieves et al., 2021; Fontanier et al., 2022; Li and Wang, 2022) suggest that this hierarchical organization of timescales may be a fundamental principle of brain organization. Despite this, analyses of single neuron timescales across humans and animal models utilizing the same data modality and fitting procedure have been lacking. This is a key consideration given the differences in temporal resolution between fMRI and single neuron activity.

Neural timescales are hypothesized to be driven by the physical properties as well as connectivity of single neurons (Soltani et al., 2021). The divergence in brain anatomy across humans, non-human primates, and rodents suggests that there could be corresponding changes in timescales across brain areas that mirror anatomical differences (Cavanagh et al., 2020). For instance, the dorsal portion of anterior cingulate cortex (ACC) is largely conserved across humans, macaques and mice as it is agranular cortex (Wise, 2008; van Heukelum et al., 2020). While the timescales of ACC neurons across primate and rodent species have not been directly compared, their shared anatomy suggests that they may be similar. By contrast, amygdala, an area which sends strong projections to ACC and other frontal cortex areas (Ghashghaei et al., 2007; Hintiryan et al., 2021; Zeisler et al., 2023), is markedly different across humans, macaques and mice – particularly in the basolateral subdivision (Chareyron et al., 2011; Zeisler et al., 2024). Whether these differences in amygdala anatomy are mirrored by differences in timescale is unknown.

To address these issues, we compared the timescale of tens of thousands of neurons within frontal cortex, amygdala, and hippocampus of mice, macaque monkeys, and humans. We identified largely consistent hierarchies across species within this amygdala-hippocampus-frontal cortex system, with key exceptions particularly in amygdala. Importantly, divergences from the common hierarchy were not related to known differences in frontal cortex cytoarchitecture.

## Results

We analyzed the firing activity of 17,458 single neurons recorded in orbitofrontal cortex (OFC), medial prefrontal cortex (mPFC), anterior cingulate cortex (ACC), amygdala and hippocampus in three species: humans, macaque monkeys, and mice (**Figure 1A**, **Table 1**, see **Methods** for detailed explanation on anatomical mapping). Multiple methods have been put forth to calculate the temporal timescales of single neurons. Timescale estimation originally relied only on the spiking activity during periods in which animals were not actively performing a task but rather sitting in restful attentiveness (Murray et al., 2014). While well-suited for analysis of data recorded from non-human primates, who often perform tasks with rigid temporal structure, such an approach is not optimal for recordings from rodents or humans, where tasks are often performed with more continuous and variable time courses. One method based on the distribution of inter-spike intervals (ISI) allows for the estimation of timescales irrespective of task events and could therefore be readily applied to the activity of neurons recorded in different species (Fontanier et al., 2022). Importantly, the estimated timescales from the approach that uses the ISI and the more standard approach (Murray et al., 2014) have been shown to be highly correlated with one another, suggesting that they are modeling the same underlying physiological properties.

**Figure 1.**
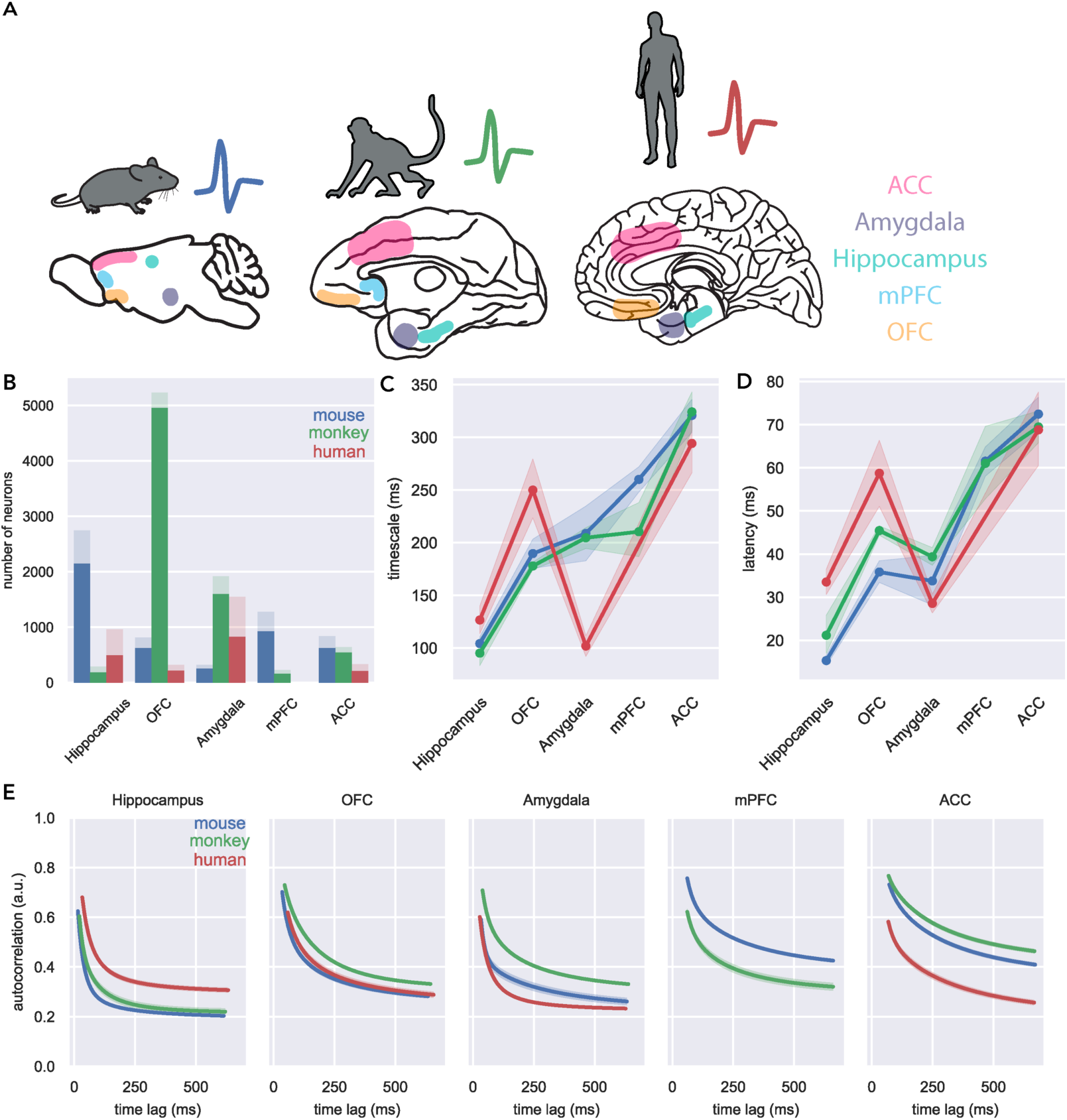
Single neuron timescale and latency hierarchies are largely conserved across species. A) Schematic representing the approximate position of brain areas from which recordings were collected in mice (left), macaques (middle) and humans (right); ACC in pink, amygdala in purple, hippocampus in cyan, mPFC in light blue, OFC in orange (see **Methods** for detailed explanation of area assignments). B) Number of neurons considered (light colors) and successfully fit (dark colors) across all brain areas. C) Average [±95% CI] timescales and D) latencies for each area in humans (red), macaques (green) and mice (blue). There was a main effect of both species (F(2) = 48.37, *p* = 1.2*10^-21^) and area (F(4) = 422.15, *p* = 0) on timescale as well as a significant interaction (F(8) = 31.53, *p* = 1.7*10^-49^). All three species had different OFC timescales (OLS regression, mouse vs monkey: t(2) = 1.961, *p* = 0.05; monkey vs human t(2) = 7.332, *p* = 2.6*10^-13^), OFC latency (mouse vs monkey: t(2) = 5.672, *p* = 1.5*10^-8^; monkey vs human t(2) = 4.788, *p* = 1.7*10^-6^), and amygdala latency (mouse vs monkey: t(2) = 2.096, *p* = 0.036; monkey vs human t(2) = 6.379, *p* = 2.1 *10^-10^). Human amygdala timescale was different from the other species (mouse vs monkey t(2) = 0.338, *p* = 0.735; monkey vs human t(2) = 13.331, *p* = 2.6*10^-39^). E) Average fits for neurons from each area and species. Lines are plotted by the mean fit parameters from successfully fit neurons, with shading indicating the standard error.

**Table 1.** Datasets used for cross-species analysis.

| First Author | Species | Brain areas | # single neurons | Number of subjects | Analysis Paper Link | Dataset link |
| --- | --- | --- | --- | --- | --- | --- |
| **Chandra-vadia**<br>(Chandravadia et al., 2020) | Human | Amygdala<br>Hippocampus | 1086<br>759 | 59 | https://doi.org/10.1038/s41597-020-0415-9 | https://doi.org/10.17605/OSF.IO/HV7JA |
| **Minxha**<br>(Minxha et al., 2020) | Human | Amygdala<br>ACC<br>Hippocampus | 460<br>329<br>203 | 13 | https://doi.org/10.1126/science.aba3313 | https://osf.io/u3kcp/ |
| **Aquino**<br>(Aquino et al., 2024) | Human | OFC | 320 | 9 | https://doi.org/10.1523/JNEUROSCI.1628-23.2024 | https://osf.io/5rv8f/ |
| **Fontanier**<br>(Fontanier et al., 2022) | Macaque | ACC | 298 | 2 | https://doi.org/10.7554/eLife.63795 | https://doi.org/10.5281/zenodo.3947745 |
| **Stoll** (Stoll and Rudebeck, 2024a) | Macaque | Amygdala<br>OFC<br>ACC | 1691<br>5230<br>343 | 2 | https://doi.org/10.1016/j.neuron.2024.03.020 | Unpublished |
| **Wirth**<br>(Wirth et al., 2017) | Macaque | Hippocampus | 290 | 2 | https://doi.org/10.1371/journal.pbio.2001045 | http://crcns.org/datasets/hc/hc-12/about-hc-11 |
| **Young/Mosher**<br>(Young et al., 2023) | Macaque | Amygdala<br>mPFC | 229<br>230 | 2 | https://doi.org/10.1016/j.neuron.2023.07.012 | https://zenodo.org/records/8164440 |
| **Steinmetz**<br>(Steinmetz et al., 2019) | Mouse | ACC<br>Amygdala<br>Hippocampus | 841<br>317<br>2745 | 9 | https://doi.org/10.1038/s4 | https://figshare.com/articles/dataset/Dataset_fro |
|  |  | mPFC | 1274 |  | 1586-019- | m_Steinmet |
|  |  | OFC | 813 |  | 1787-x | z_et_al_201 |
|  |  |  |  |  |  | 9/9598406 |

To estimate the intrinsic timescale of each of the neurons we analyzed, we employed this latter ISI-based method from Fontanier, Sarazin and colleagues (Fontanier et al., 2022). In brief, we fit an exponential decay function to the descending portion of the distribution of ISIs according to the following equation:

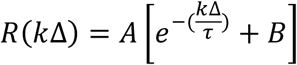

where *k*Δ is the time lag between bins, τ represents the neuron’s *timescale*, describing the rate of decay of the autocorrelation distribution, and A and B are free parameters which together describe the peak and asymptotic limit of the autocorrelation. We also extracted the time bin at which the autocorrelation begins decreasing, which we refer to as *latency*. Only neurons for which the model fit well (R^2^ > 0.5) and exhibiting timescales within the range 10 < τ < 1000 were included for further analysis (n=13,755 single neurons, 78.7% success; **Figure 1B**).

Mice, macaques, and humans exhibited largely similar hierarchies of single neuron timescales and latencies across areas (**Figure 1C, D**). Across all three species, ACC neurons showed the longest timescales and latencies while hippocampus neurons showed the shortest timescales. Other areas were characterized by greater across-species variability. Notably, human amygdala neurons had a significantly shorter timescale compared to macaques and mice (**Figure 1E**; middle). All three species also had different amygdala latencies. Furthermore, the timescales of single neurons in OFC were statistically distinguishable in all three species, with humans exhibiting the longest timescale. Taken together, we found numerous similarities between the timescale and latency hierarchies across these three species, with a notable exception for OFC and amygdala neurons in humans.

While the above analyses focused on the average timescales for each area and species, timescales were not necessarily uniform within a given area. Indeed, we found large variability in the distributions of timescale and latency within each area and across all three species (**Figure 2 A, C-D)**. In fact, both timescale and latency distributions across most considered areas and species had unequal variances. This within-area variation is particularly prominent in ACC, where mice, macaques, and humans all had a bimodal distribution of timescale: a small population of neurons with short timescale and a larger population of neurons with longer timescales. Such consistent heterogeneity across species within an area potentially provides insight into the organizing role that timescales might play in the temporal processing of information. Indeed, it could be the case that the distribution of timescales within an area is more critical for supporting that area’s function than the average timescale.

**Figure 2:**
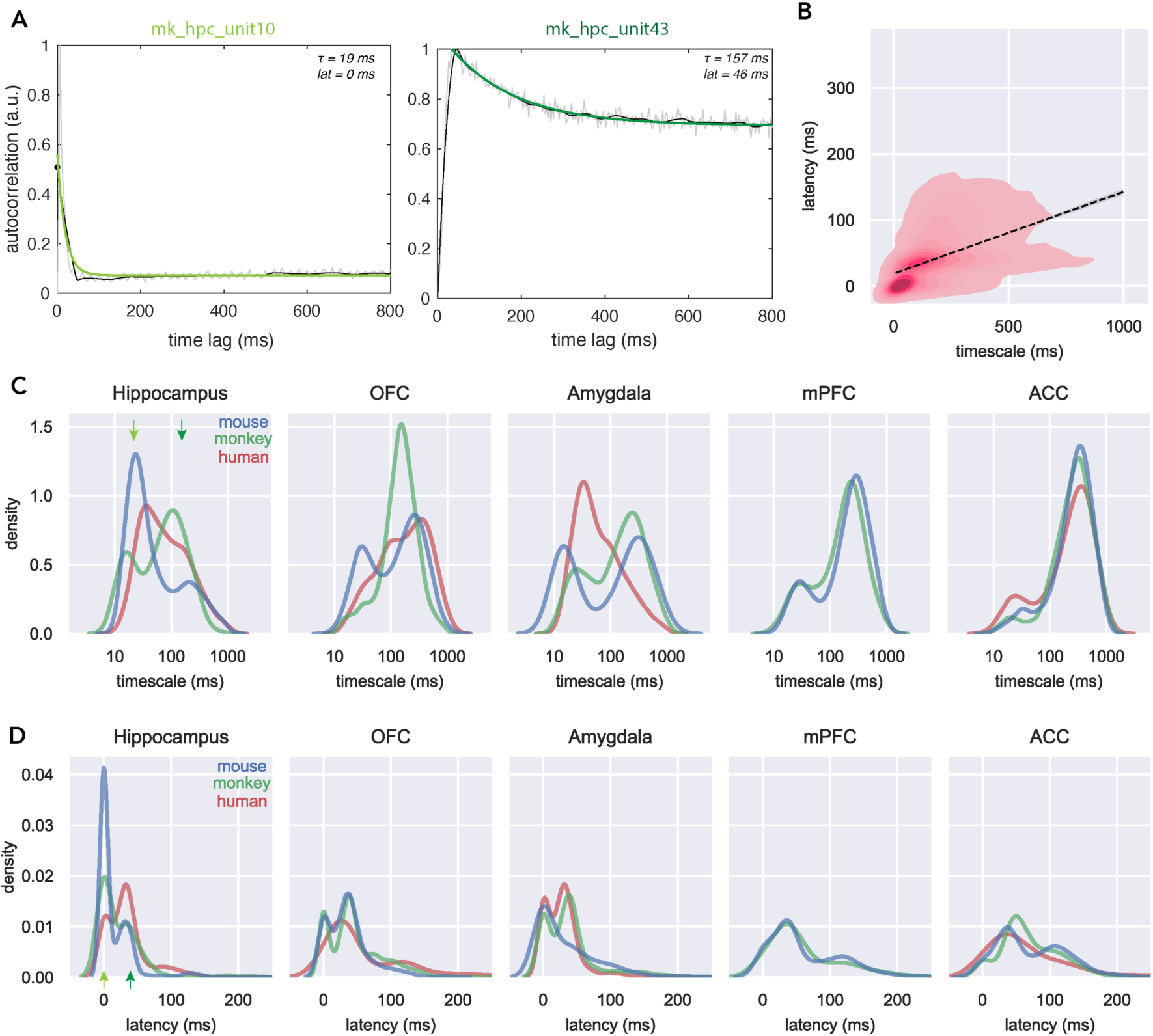
Distributions of timescale and latency for individual neurons. A) Two example single-neuron fits from macaque hippocampus with relatively short (left; light green) and long (right; dark green) timescales. Grey traces indicate normalized ISI distribution, the black line is the smoothed autocorrelation, and the green lines are the fit autocorrelation function, specified by timescale (*τ*) and latency (*lat*) shown in the top right of each panel. B) Density plot of timescale and latency, with regression line overlaid. Timescale and latency are correlated with each other (OLS regression: R^2^ = 0.501, *p* < 0.001). C) Density histograms showing distribution of timescales across neurons recorded within each brain area. The timescales of neurons from OFC, amygdala, and mPFC all have unequal variance across species (Brown-Forsythe test: p < 0.004). Green arrows indicate the position of the example cells shown in panel A. D) Same as C for latencies (bottom). All areas with the exception of mPFC had unequal variance regarding latencies (p < 10^-6^).

Across the different areas and species, there was also significant variability in the latency measure that was fitted to each neuron’s spiking activity (**Figure 1D**). Here ACC and mPFC had the longest latency whereas hippocampus generally had the shortest. Further, within each area there was also a high degree of variability in the latency (**Figure 2D**). Importantly, while timescale and latency exhibit some degree of correlation (**Figure 2B**), each explains only half of the variance in the other. This point highlights that latency is a factor that is partially separable from the timescale and likely contributes to how information is temporally processed within a brain area.

One potential issue with fitting exponential decay functions to the activity of single neurons is that they might be influenced solely by the firing rates of the neurons. Thus, we next checked that our timescale and latency estimates reflected distinct neuronal functional properties, independent from the influence of firing rate. To that end, we built regression models to predict timescales based on spiking statistics, areas and species (**Supplemental Figure 1**). We found that the number of spikes entered into the analysis only had a significant influence on a neurons’ timescale while the number of spikes, mean firing rate, and Fano factor all influenced the neural latencies. These parameters, however, offered little additional explanatory power compared to species and area, explaining significantly less variance than area and species alone. Thus, the neural timescale and latency that we reported above (**Figures 1, 2**) represent features of neuronal dynamics that are distinct from classical spiking statistics.

### The relationship between cytoarchitecture and timescales within ventral frontal cortex

As previously noted, ACC in humans, macaques, and mice is agranular cortex (van Heukelum et al., 2020). By contrast, OFC in macaques (**Figure 3A**) and humans is composed of granular, dysgranular, and agranular cortex (Carmichael and Price, 1994; Wise, 2008). OFC in mice, by contrast, is entirely agranular. One possibility is that these differences in granularity might underlie the timescale and latency differences we observed in OFC across species (**Figure 1**). To assess this, we concentrated our analysis on recordings from 12,043 single neurons in macaque ventral frontal cortex, of which 5,230 were included in the previous analyses of macaque OFC in **Figures 1** and **2**. This dataset included the activity of single neurons recorded using 157-channel semi-chronic microdrives in two monkeys, where each electrode trajectory was precisely reconstructed (Stoll and Rudebeck, 2024b, 2024c). Using standard histological techniques, the ventral frontal cortex was subdivided into 8 cytoarchitectonic areas based on analysis of Nissl-, SMI-32- and calbindin-stained tissue series (**Figure 3A**). This allowed us to pinpoint the precise location of recorded neurons according to previous parcellations of the ventral frontal cortex of macaques (Carmichael and Price, 1994; Rapan et al., 2023). Thus, we could compare the timescales of neurons from neighboring subdivisions with distinct cortical granularity; fully granular areas 11m/l, 12l, 12m, and 12o, dysgranular areas 12r, 13m, and 13l areas, and agranular insula (AI; the number of neurons in each subdivision is shown in **Figure 3B** and **Table 2**).

**Figure 3.**
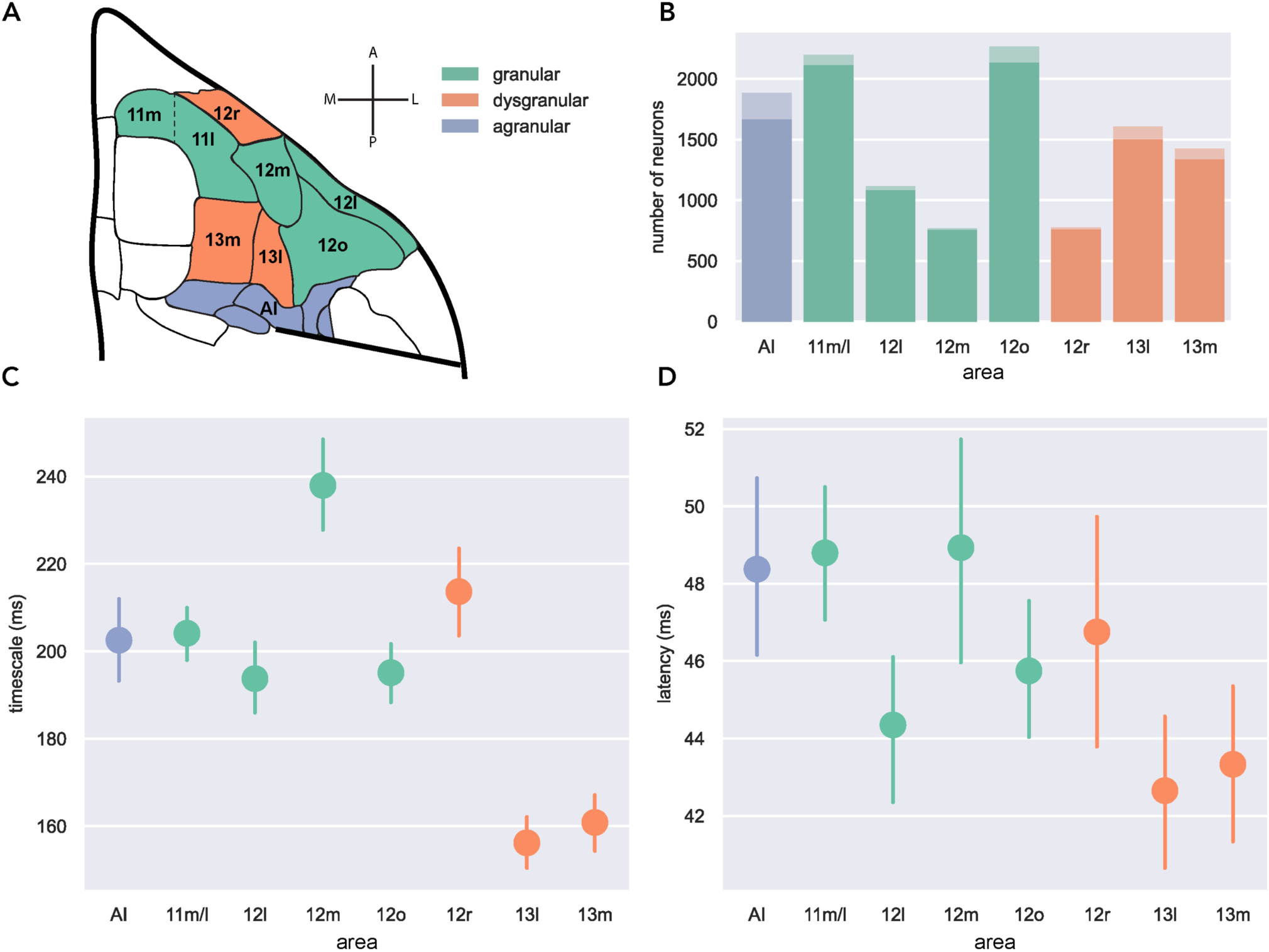
Timescales and latencies within macaque ventral frontal cortex are not purely driven by cortical granularity. A) Schematic representations of the orbital surface of macaque’s brain showing the relative location and cytoarchitectonic subdivisions of the ventral frontal cortex included in this analysis. B) Number of recorded neurons (lighter bars) and number of neurons successfully fit (darker bars) across areas. C) Average (+/-95% confidence intervals) timescale of single neurons in granular (green), dysgranular (orange) and agranular (blue) subdivisions of ventral frontal cortex. There is a significant main effect of granularity (OLS regression, dysgranular < granular: t(2) = 10.58, p = 4.7*10^-26^; dysgranular < agranular: t(2) = 7.34, p = 2.36*10^-13^; agranular ≈ granular: t(2) = .19, p = 0.85); although a model which includes area and granularity is no better at predicting timescale than one which includes area alone (F(2) = 1.36,

**Table 2.**
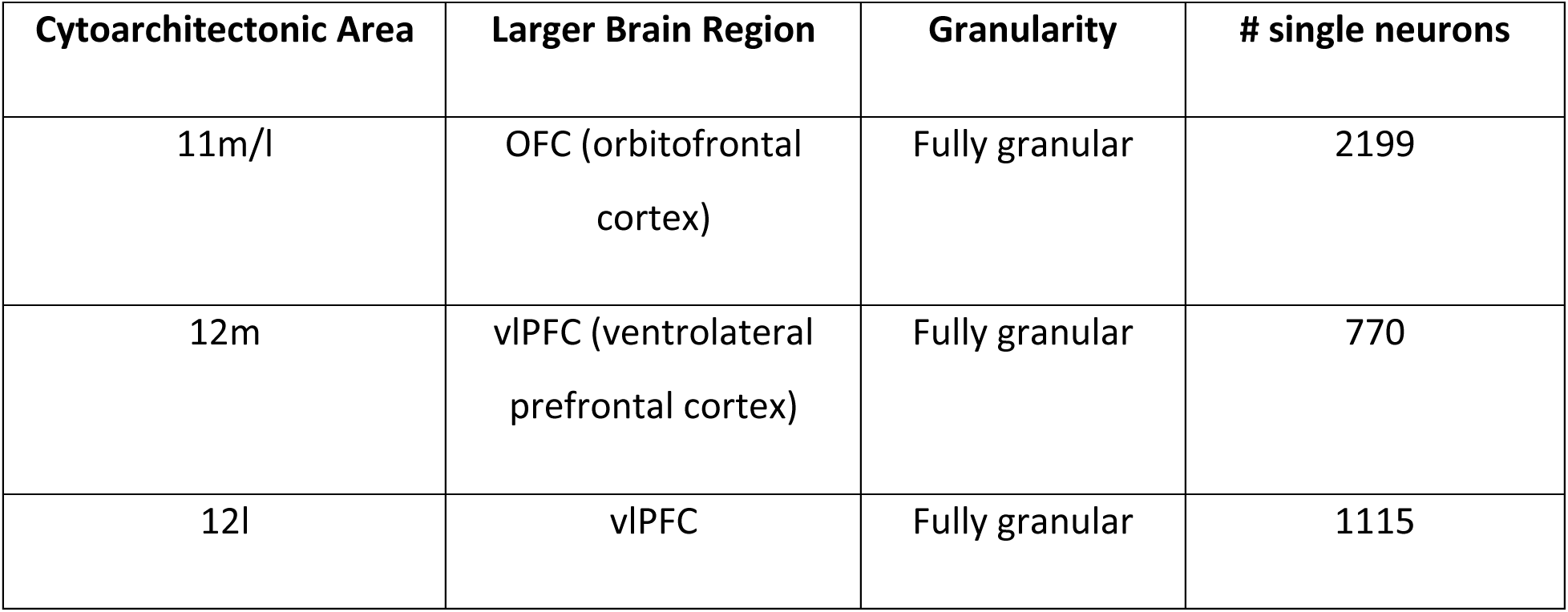
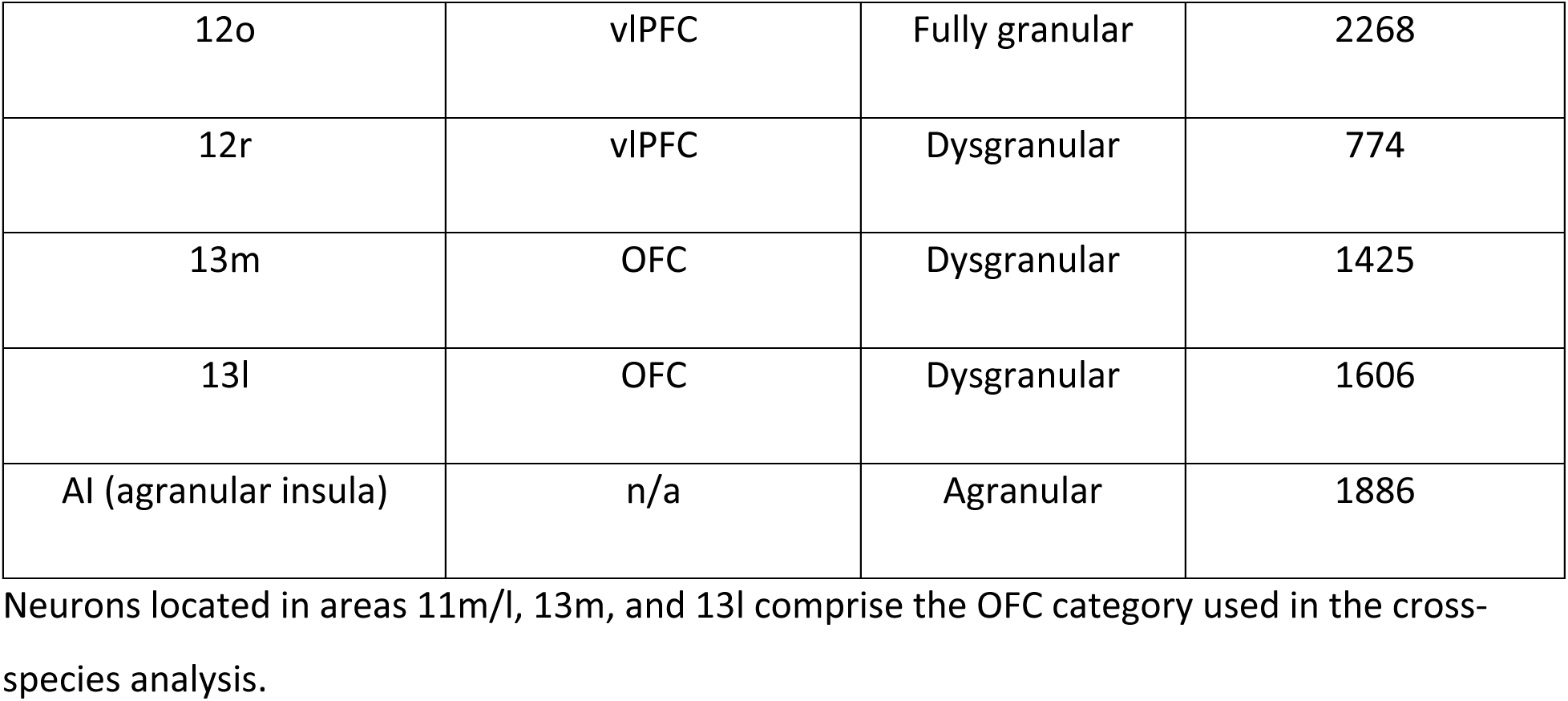
Data used for within-frontal cortex analysis.

We found that timescale variability across macaque ventral frontal cortex could not be fully explained by differences in granularity (**Figure 3C**, left). While there was a main effect of granularity, such that neurons in granular areas tended to exhibit longer timescales than those in dysgranular areas, hierarchical model comparison indicated that granularity did not significantly improve a model predicting neural timescales based on area alone. Notably, adjacent areas 12m and 12r were most different from other granular and dysgranular areas, respectively as they had relatively longer timescales. Neuronal latencies were found to have a different relationship to granularity (**Figure 3D**, right). Specifically, granular and agranular areas demonstrated longer latencies than dysgranular – and a model predicting latency with granularity performed better than a model containing area alone. Thus, the presence or absence of an internal granule cell layer does not appear to be a strong predictor of the timescale of single neurons in an area. Granularity, however, does significantly affect neuronal latency, which may be related to variation in the cortical microstructure (Gao et al., 2020).

Finally, we investigated whether single neuron timescales and latencies could be related to the spatial location of neurons on the ventral surface of the frontal lobe. Across frontal cortex there are both antero-posterior (AP) and medio-lateral (ML) gradients in the connections from other parts of the brain, especially the amygdala (Ghashghaei et al., 2007) and hippocampus (Aggleton et al., 2015). We hypothesized that such differences in long-range inputs could drive differences in neural timescales. To this end, we assessed whether the AP and ML coordinates of each recorded neuron influenced neuronal timescales and latencies. In both animals, there were main effects of the AP position (F(1) = 20.16, p = 7.3*10^-6^), ML position (F(1) = 26.97, p = 2.1*10^-7^), and a significant interaction (F(1) = 47.12, p = 7.5*10^-12^), such that timescales were longest in neurons recorded in posterior/lateral and anterior/medial regions. Thus, the variation in timescales in ventral frontal cortex could be influenced by the patterns of projections into the frontal cortex from limbic (Cavada et al., 2000; Ghashghaei et al., 2007) and/or sensory systems (Carmichael and Price, 1996).

## Discussion

Here, we computed the timescales of single neurons recorded in parts of the fronto-limbic system in humans, macaques, and mice. We found that across the different species, single-neuron timescales followed a similar hierarchical organization; hippocampus had the shortest timescale whereas ACC had the longest. Further, while timescales in hippocampus, ACC, and medial frontal cortex were highly consistent across species, the timescales of amygdala and OFC neurons were more variable. We further found that differences in timescales across macaque ventral frontal cortex followed specific spatial patterns but were independent of the underlying variation in cytoarchitecture. Taken together, we report that hierarchical organization of brain systems is common between three mammalian species frequently studied in neuroscience, despite differences in some areas.

### Timescales in subcortical areas

One of the most striking results from these data was the large divergence in amygdala timescale across the species that we studied (**Figure 1**). We observed that mice had the longest amygdala timescales while human amygdala timescales were the shortest. It is important to note that within these mammals, large anatomical differences between the amygdala of different species exist (Chareyron et al., 2011). Such differences in amygdala structure are most dramatic within the basolateral subdivision, the target of the recordings considered in our study. Other parts of amygdala like the central nucleus, by contrast, are more similar across primates and rodents and it is possible that timescales could be more conserved in these nuclei. Why this difference in basolateral timescales exists between the species is unknown, but work by Nougaret et al. reported gradients of timescales within the basal ganglia, a circuit with well-defined input and output structure (Nougaret et al., 2021). This then suggests that an area’s outputs might contribute to determining its timescale. Thus it follows that the basolateral amygdala, which primarily projects to frontal cortex, might have longer timescales than central nucleus, which primarily projects to brainstem and hypothalamus (Ghashghaei et al., 2007). Future experimental work and analyses could intentionally characterize neurons from separate amygdala subdivisions to determine whether anatomical and functional organization might also extend to differences in timescales.

The macaque amygdala has a higher glia to neuron ratio, more sparse distribution of neurons, and greater density of neuropil compared to rats (Chareyron et al., 2011). This difference is even more stark when comparing to humans, where neurons in amygdala are even more sparsely distributed than in macaques. Chareyron and colleagues speculate that these specifications predispose the primate amygdala to a preferential role in integrating information coming from a dramatically expanded cortex compared to rats. The basal and lateral nuclei receive much of the input from the cortex and send intra-amygdala projections to other nuclei (Freese and Amaral, 2009), placing them in a key position to perform such an integrative role. In sensory areas, short timescales are hypothesized to be critical to quickly process and update incoming information. Such shorter timescales in amygdala in humans and macaques compared to mice could therefore predispose neurons in these species to be able to quickly integrate information from the cortex which would in return enable the amygdala to drive learning in areas with longer timescales, such as ACC (Stolyarova et al., 2019). That the primate amygdala serves a more integrative role is further supported by recent insights about the organization of projections from this area. Specifically, basolateral amygdala neurons in macaques exhibit highly-organized patterns of collateral projections to the frontal lobe (Zeisler et al., 2023), while those in mouse amygdala project more broadly in a less specific fashion (Zeisler et al., 2024). While technically challenging, comparing the timescale of individual neurons whose anatomical connections are known could bring important insight in our understanding of how neuronal branching is related to temporal dynamics.

Finally, the cross-species variability in amygdala timescales stands in stark contrast to hippocampus, where timescales were shorter than other brain areas measured here and highly similar across species. The hippocampus exhibits much less differentiation across mammalian species (Manns and Eichenbaum, 2006), with little about its anatomy and possibly the computation it subserves changing throughout the course of evolution (Clark and Squire, 2013). Thus, our finding that hippocampal timescales are similar across species appears to correlate with such conservation. It’s also striking that hippocampus had such short timescales given that previous reports refer to primary sensory cortices as the prototype of an area with a short timescale (Murray et al., 2014; Manea et al., 2022). Perhaps such similarity is driven by the fact that most hippocampal inputs arise from the entorhinal cortex (Witter et al., 2000), in the same way that primary sensory cortex inputs are predominantly from their respective sensory organs.

### Anatomy, granularity, commonality

It remains unclear what exactly about a brain area or a neuron sets its intrinsic timescale. Possible contributing factors include region-wide variation in receptor expression, genetic identity, intrinsic recurrence, or inputs from distant areas (Cavanagh et al., 2020; Froudist-Walsh et al., 2021; Rapan et al., 2023). Indeed, we identified differences across monkey cytoarchitectonic ventral frontal subdivisions (**Figure 3**). Such variation could also be more significantly driven by network connectivity. Carmichael and Price have described two largely non-overlapping networks within these regions (Carmichael and Price, 1996): the orbital or so-called sensory network, and the medial or viscero-motor network. Recent functional connectivity work (Stoll and Rudebeck, 2024b) has also explored information flow between these regions and revealed that both areas 13m and 13l lack strong functional connectivity with other OFC subregions during this particular decision-making paradigm. This low intra-frontal connectivity might be influencing the relatively low timescale in these two subdivisions of area 13. By contrast, Stoll and Rudebeck show that the nearby area 12o is very highly interconnected with other prefrontal regions; 12o is also more likely to receive information from many other areas than to send it (Stoll and Rudebeck, 2024b). These results suggest that the highly integrative role of area 12o might contribute to its relatively long timescale. Further evidence for this relationship between connectivity and timescale could come from future experiments targeting the visual system, with its well-documented organization of feed-forward and feedback connections (Vanni et al., 2020), as one would expect shorter timescales in earlier visual areas like V1 and longer timescales further up the visual hierarchy in V4 or parietal cortex.

Within OFC, we identified spatial gradients of timescale that could not be accounted for by cortical granularity alone (**Figure 3**). Rather, we found that timescales varied in a patchy organization, with no clear pattern along a single anatomical axis. What drives this variation is unclear, although it could be related to the projections coming into the ventral frontal cortex. Inputs from the amygdala and hippocampus exhibit a sparse distribution in ventral frontal cortex (Aggleton et al., 2015). Hippocampal inputs to OFC preferentially target more medial and posterior areas, while amygdala inputs project more strongly to lateral areas (Ghashghaei et al., 2007). Perhaps this difference in long-range innervation could drive differences in timescale across the ventral surface of macaques. Within this view, the short timescale of hippocampal neurons might entrain the neurons to which they project, although we note that the precise effect of the presynaptic neurons’ timescale on its downstream targets has yet to be assessed.

While often overlooked in analyses of timescales, we found that the presence or absence of granule cells in layer IV modulated the latency of the exponential decay (**Figure 3B**). Prior work by Fontanier, Sarazin and colleagues found that timescale and latency tend to vary together and biophysical modeling revealed that the latency of a particular neuron was closely related to potassium and other cationic currents (Fontanier et al., 2022). By contrast, we found that variation in latency was related to granularity across macaque ventral frontal cortex, whereas such a relationship was not present for timescale. Such a difference supports the idea that the latency measure is distinct from timescale. Indeed, while there is consensus that a longer timescale indicates a longer time interval over which a neuron might integrate information (Soltani et al., 2021), a functional role for latency remains to be described. Further work both *in vivo* and *in silico* is needed to fully understand how variations in latency might alter computation.

Our analyses revealed a diversity of within-area timescales that were in some cases consistent across species (**Figure 2**). The extent of these within-area differences has not been previously reported. Such heterogeneity is hypothesized to be important for optimal functioning both in brains and in artificial neural networks (Cavanagh et al., 2020; Stern et al., 2023). Artificial neural network (ANN) modelling of prefrontal activity during a working memory task suggests that timescale diversity is an adaptation to facilitate task performance; the network had a largely homogenous timescale distribution before training which widened significantly after repeated training iterations. Using brain-inspired spiking ANNs, Perez-Nieves and colleagues successfully determined that heterogeneity is a feature of brain that is critical to enable problem-solving (Perez-Nieves et al., 2021). ANN modeling therefore highlights the computational significance of diverse timescales as well as the need for such heterogeneity to both accurately model the brain during task performance.

While the timescales of neurons varied within an area we found that they were not uniform but often exhibited a bimodal distribution (**Figure 2**). Importantly, a recent analysis of intrinsic timescales in the macaque frontal eye field (FEF; Soyuhos et al., 2024), an area critical for directing attention and coordinating eye movements, also identified bimodal distributions of timescales. By characterizing these two populations using their functional properties during a passive viewing task, they reported that short timescale FEF neurons primarily processed rapid visual features of the task, while long timescale FEF neurons were related to sustaining attention. Although a bimodal distribution was not directly reported, other studies have also identified a relationship between variation in timescale and functional properties (Cavanagh et al., 2020; Spitmaan et al., 2020; Fontanier et al., 2022). Since we did not characterize the functional properties of the neurons included in our analysis, it remains an open question whether bimodality in other brain areas is driven by related features. Further, while nearly all timescale distributions considered in this analysis were bimodal, some species had uniquely unimodal distributions: for example, macaque OFC or human amygdala. Whether this effect is a result of the precise areas sampled or the task performed by subjects, understanding to what extent bimodal distributions of timescales are ubiquitous across brain areas and behaviors is an exciting next direction both for neuroscience and for computational modeling. Future research should strive to further elucidate how the structure of variation of within-area timescales across the cortex and subcortical structures contributes to the computational roles of those areas in behavior.

Taken together, our data show that there are consistent timescale and latency hierarchies across three commonly utilized model species: humans, macaque monkeys, and mice. While we were able to reveal clear differences, particularly in OFC and amygdala, the high degree of consistency across species suggests that a hierarchy of timescales might be a broad organizing principle of mammalian brains. The implementation of such principles in neural networks could provide improved sensitivity to both short- and long-term variation in environmental inputs.

## Methods

Analyses were performed using MATLAB R2021a (MathWorks) and Python 3.9. All figures made in Python used Seaborn (Waskom, 2021) and Matplotlib (Hunter, 2007) packages. Other specific packages used will be cited in the methods below. Analysis code is available on GitHub: https://github.com/RudebeckLab/timescales.

### Neurophysiological datasets

Here, we analyzed the activity of single neurons recorded in humans, macaques and mice across both open-source and unpublished datasets. **Table 1** illustrates the sources, primary authors, and web links to these datasets as well as the papers in which they were originally presented, if applicable. Unpublished datasets are indicated as such.

### Operational Definitions of Brain Regions

For the sake of clearer comparisons, both within and across species, we combined recordings into larger brain regions.

From mice (Steinmetz et al., 2019), hippocampal subfields CA1, CA2, and Dentate Gyrus were all combined into the hippocampus category; Anterior cingulate area (ACA) was assigned to ACC; Basolateral amygdala (BLA) to amygdala; Infralimbic area (ILA) and prelimbic cortex (PL) to the mPFC category (Wise, 2008; Wallis et al., 2017); Orbital cortex (ORB) to OFC.

From macaque monkeys, amygdala recordings primarily targeted BLA, though precise anatomical placement within amygdala was not formally assessed; OFC recordings were taken from Brodmann’s areas 11 and 13; subcallosal ACC (Walker’s area 25) was assigned to the mPFC category (Wise, 2008; Wallis et al., 2017); hippocampal recordings (Wirth et al., 2017) were taken from across the extent of the hippocampus both in the antero-posterior and medio-lateral planes; mid-cingulate cortex (MCC) recordings (Fontanier et al., 2022; Stoll and Rudebeck, 2024a) were assigned to the ACC category, as this area has been referred both as anterior MCC or dorsal ACC (Procyk et al., 2016).

From humans (Chandravadia et al., 2020; Minxha et al., 2020; Aquino et al., 2024), all recordings were performed during seizure monitoring in cases of drug-resistant epilepsy; recording targets were exclusively chosen according to clinical criteria. Amygdala recordings preferentially targeted the BLA; ACC recordings were from Brodmann areas 24 and 32; hippocampal recordings covered the extent of the hippocampus both in the A/P and M/L planes; OFC recordings were taken primarily from Brodmann’s areas 14, 13, and 10.

### Macaque ventral frontal cortex dataset

Neural recordings were collected from two male rhesus macaques performing one of two behavioral tasks: either an instrumental preference task described in Stoll and Rudebeck (Stoll and Rudebeck, 2024a), or a learning task in which animals associated novel visual stimuli with different reward probabilities which changed over the course of the session (unpublished).

Recording procedures are described in more detail in Stoll and Rudebeck (Stoll and Rudebeck, 2024a). Briefly, animals were implanted with 157-channel semi-chronic recording microdrives (Gray Matter Research). During recordings, animals were seated in primate chairs while electrophysiological signals were recorded using the Plexon Omniplex system. Putative single neurons were determined using MountainSort (Chung et al., 2017) and manually curated based on cluster isolation metrics.

Electrodes were moved each day, and their cumulative adjustments tallied to determine their precise location in the brain. CT images collected throughout the recordings and histological assessment after perfusion were also used to verify these locations. Specifically, following standard perfusion with 4% formaldehyde (Electron Microscopy Sciences) in phosphate-buffered saline (PBS, Fisher), brains were extracted and post-fixed for 24-48 hours. Brains were then cryoprotected in increasing sucrose solutions (10%, 20%, 30% sucrose in PBS) before being shipped to FD NeuroTechnologies, Inc for processing, the details of which are also described in detail(Stoll and Rudebeck, 2024b). Brains were further cryoprotected, frozen in isopentane, and sectioned at 50um on a sliding microtome. Then equally spaced series of sections were prepared for either Nissl staining (one series) or immunohistochemistry (three series). Immunohistochemistry was performed for calbindin and SMI-32 in order to delineate boundaries between cortical areas.

Boundaries between cortical areas were determined according to features described previously (Carmichael and Price, 1994; Rapan et al., 2023; Stoll and Rudebeck, 2024b). In the most anterior sections, areas 11m/l, 12m, and 12r were separated primarily based on SMI-32 staining (11m and 11l were not consistently distinguishable in these animals and so have been left grouped together). 11m/l is denser with cells than the more medial 13b, and it is denser with superficial neurites in layer II/III than the more lateral 12m. Then, in Nissl-stained sections, the 12r and 12m can be separated based on the dysgranular layer IV in the former compared to the fully granular later IV in the latter. More posteriorly, area 13l was distinguished from 13m by the presence of vertical processes in calbindin-stained sections, particularly in layers III and V. Area 12o was defined by its stark lack of neurite staining in SMI-32-labeled sections compared to the more medial 13m. 12l was distinguished from 12o by significantly increased density of SMI-32-labeled cells. Finally, agranular insular areas were distinguished from area 13l on the medial side and 12o on the lateral by the distinct lack of a granular layer IV; instead, significant vertical processes were observed alongside a dense and thick layer IV in SMI-32-stained sections.

### Timescale estimation using inter-spike interval-based approach

Single-neuron timescales were estimated using an inter-spike interval-based (ISI) method (Fontanier et al., 2022). Fitting was performed using MATLAB 2021a. Briefly, the time difference between every spike in a neuron’s time series and its following 100 spikes was calculated. The lagged differences were then binned at 3.33ms from 0 to 1000ms. This distribution was smoothed, after which we fitted the following equation:

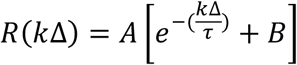

where *k*Δ is the time lag between bins, τ represents the neuron’s *timescale*, describing the rate of decay of the autocorrelation distribution, and A and B are free parameters which together describe the peak and asymptotic limit of the autocorrelation. We also extracted the time bin at which the autocorrelation begins decreasing, which we refer to as *latency*. As before, fitting was performed using only the decreasing portion of the autocorrelation distribution (after the latency).

Post-processing and plotting was performed in Python 3.9 using the following packages: NumPy, Pandas, Matplotlib, Seaborn, SciPy, and statsmodels. Only neurons for which the model fit well (R^2^ > 0.5) and exhibiting timescales within the range 10 < τ < 1000 were included for further analysis. To assess the effects of species and brain region on neuronal timescales and latencies, we first used an ordinary least squares regression (OLS; statsmodels *ols* function). We fitted the following linear model to describe a neuron’s timescale and latency in terms of the species and brain region from which it was recorded:

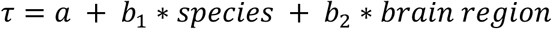

Post-hoc tests were performed within brain regions to compare species directly with FDR corrections for multiple comparisons. Timescale and latency distributions within each brain area were compared using a Brown-Forsythe test for equality of variances.

### Firing rate analyses

To ensure our timescale and latency estimates were distinct from differences in other firing rate statistics, we calculated the following measures for each neuron:

- Number of spikes (*n spikes*): number of spikes in timeseries
- Mean firing rate (*mean FR*): number of spikes divided by total length of timeseries
- Normalized FR (*norm FR*): spikes were binned at 500ms, and averaged with a sliding window of 1 second (2 bins); this firing rate was then z-scored using the mean firing rate above
- Fano: *norm FR* divided by the variance of *norm FR*
- Proportion of bursts (*prop burst*): proportion of spikes included in a burst, defined as the top 0.5 percentile of the ISI distribution(Ko et al., 2012)
- Proportion of pause (*prop pause*): proportion of spikes included in a pause, defined as the bottom 0.5 percentile of the ISI distribution(Ko et al., 2012)

All parameters were z-scored within each species and brain region. Then, we performed OLS regressions predicting both timescale and latency using each z-scored parameter individually alongside species and brain region. We extracted the percent explained variance for each parameter and computed p-values for each. Models were directly compared using ANOVA (statsmodels *anova_lm*).

### Analysis of single neuron timescales in macaque ventral frontal cortex

To understand whether timescales were influenced by cortical granularity, we analyzed an extended dataset containing single unit activity recorded from multiple subdivisions of the ventral frontal cortex in monkeys, including the previously defined OFC neurons (**Table 2**) (Stoll and Rudebeck, 2024b). We then used hierarchical model comparison to determine the relative contributions of both cortical granularity and area. For both timescale and latency, two OLS model were built, using either the specific area or the granularity as predictors. We then compared the fit of those models using *anova_lm* to determine whether the differences in degrees of freedom resulted in a better fit. Neurons’ position in 3D space were also computed using the recording depth, MRI and CT scans as well as using histological confirmation. This allowed us to assess the relationship between spatial position and timescale/latency. 3D coordinates were projected into 2D space; because our AP and DV coordinates were highly correlated with one another (more posterior neurons are more ventral), we focused our analyses only on the A/P and M/L planes. OLS regression models were built to explain the extracted timescales and latencies using the AP and ML coordinates (including the interaction term).

## Author contributions

ZRZ, FMS, and PHR conceived the study. ZRZ and FMS performed the analyses. FMS collected the macaque ventral frontal cortex dataset, and FMS and ML localized the macaque ventral frontal cortex recordings to cytoarchitectonic regions. UR provided access to the human datasets and key feedback on analysis approaches throughout. ZRZ, FMS, and PHR wrote the paper. All authors edited and approved the final version of the paper.

## Acknowledgements

The authors thank the individuals who publicly provided their data for our use in these analyses: Tomas Aquino, Nand Chandravadia, and Juri Minxha; Vincent Fontanier and Emmanuel Procyk; Sylvia Wirth and Jean-Rene Duhamel; Megan Young and Clayton Mosher; and Nick Steinmetz. ZRZ, ML, FMS and PHR are supported by grants from NIMH and the BRAIN Initiative (R01MH110822, R01MH132064). UR is supported by the BRAIN Initiative (U01NS117839).

## Supplemental figures

**Supplemental Figure 1.**
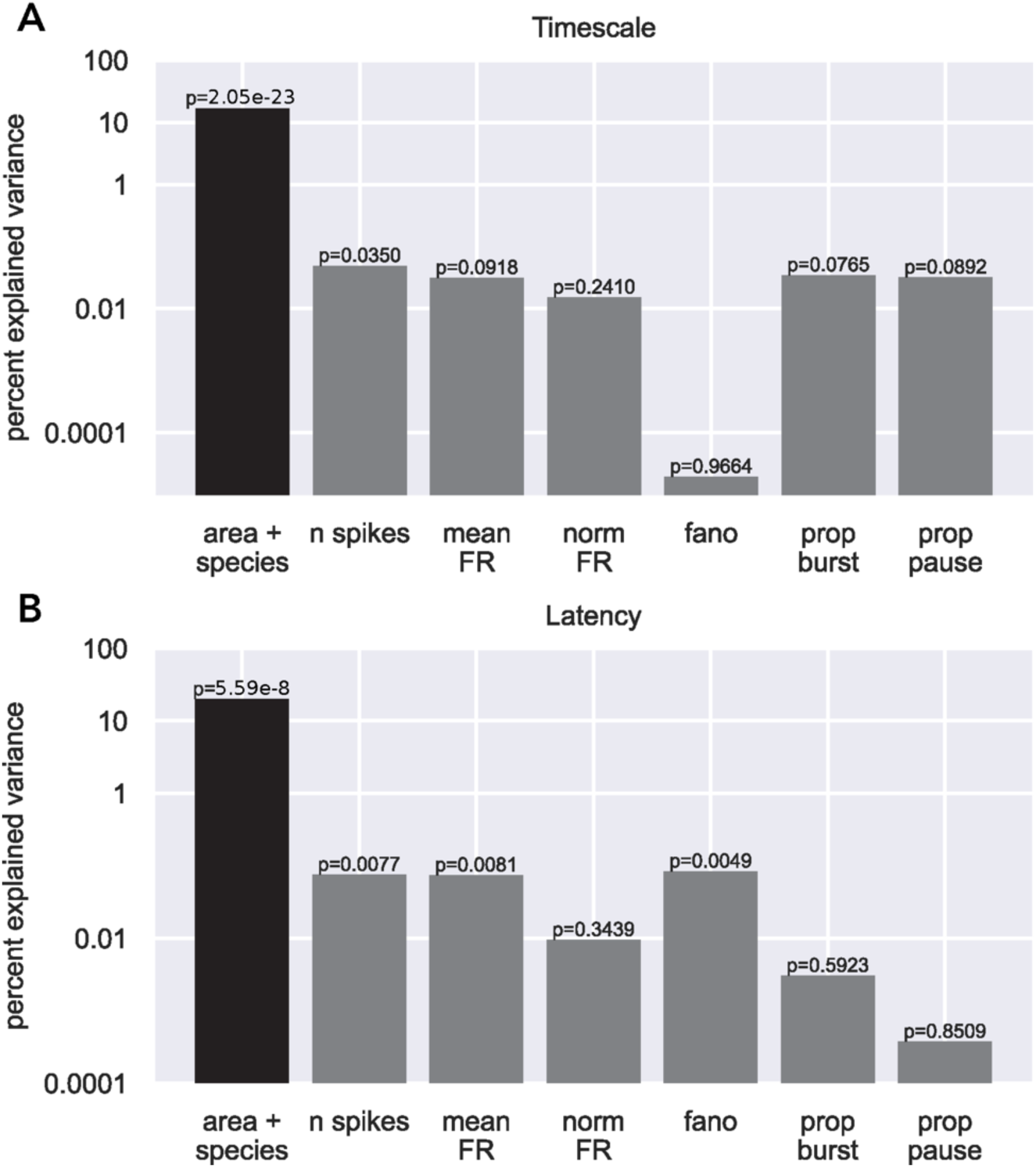
Latency and timescale are weakly related to spiking statistics. A) Percent explained variance (log scale) of timescale using a linear model containing area and species (black), and additional variance explained when including each firing rate statistic (grey). While the number of spikes was a significant predictor of timescale, the variance explained by this parameter was nearly two orders of magnitude less than the variance explained by area and species. P-values are also shown for each predictor. B) Same as A for latency. Again, while the number of spikes, mean firing rate, and Fano factor were significant predictors of latency, together, they explain less than 1% of the total variance in latency.

